# No support for white matter microstructural properties and connectivity differences in the combined and inattentive adhd presentations

**DOI:** 10.1101/2020.12.22.423936

**Authors:** Jacqueline F. Saad, Kristi R. Griffiths, Michael R. Kohn, Taylor A Braund, Simon Clarke, Leanne M. Williams, Mayuresh S. Korgaonkar

**Affiliations:** The Brain Dynamics Centre, Westmead Institute for Medical Research, The University of Sydney, Sydney, Australia; Discipline of Psychiatry, Western Clinical School, The University of Sydney, Sydney, Australia; Centre for Research into Adolescents’ Health, Department of Adolescent and Young Adult Medicine, Westmead Hospital, Sydney, NSW, Australia; Department of Psychiatry and Behavioral Sciences, Stanford University, Stanford, California, USA; Sierra-Pacific Mental Illness Research, Education, and Clinical Center (MIRECC), VA Palo Alto Health Care System, Palo Alto, California, USA

## Abstract

Evidence from functional neuroimaging studies support neural differences between the Attention Deficit Hyperactivity Disorder (ADHD) presentation types. It remains unclear if these neural deficits also manifest at the structural level. We have previously shown that the ADHD combined, and ADHD inattentive types demonstrate differences in graph properties of structural covariance suggesting an underlying difference in neuroanatomical organization. The goal of this study was to examine and validate white matter brain organization between the two subtypes using both scalar and connectivity measures of brain white matter. We used both tract-based spatial statistical (TBSS) and tractography analyses with network-based Statistics (NBS) and graph-theoretical analyses in a cohort of 35 ADHD participants (aged 8-17 years) defined using DSM-IV criteria as combined (ADHD-C) type (n=19) or as predominantly inattentive (ADHD-I) type (n=16), and 28 matched neurotypical controls. We performed TBSS analyses on scalar measures of fractional anisotropy (FA), mean (MD), radial (RD), and axial (AD) diffusivity to assess differences in WM between ADHD types and controls. NBS and graph theoretical analysis of whole brain inter-regional tractography examined connectomic differences and brain network organization, respectively. None of the scalar measures significantly differed between ADHD types or relative to controls. Similarly, there were no tractography connectivity differences between the two subtypes and relative to controls using NBS. Global and regional graph measures were also similar between the groups. A single significant finding was observed for nodal degree between the ADHD-C and controls, in the right insula (corrected p=.029). Our result of no white matter differences between the subtypes is consistent with most previous findings. These findings together might suggest that the white matter structural architecture is largely similar between the DSM-based ADHD presentations.

## Introduction

Characterized by clinical deficits in attention, hyperactivity, and impulsivity, attention deficit hyperactivity disorder (ADHD) is a highly prevalent neurodevelopmental condition with an estimated global prevalence of 3.4% of children worldwide (1). Presentation types (i.e. subtypes) of ADHD are categorized as either predominantly inattentive (ADHD-I), predominantly hyperactive-impulsive (ADHD-HI) or combined (ADHD-C) (2). Cognitive and behavioral differences between clinical presentations have been well documented across the neuropsychological literature (3). Furthermore, the knowledge gap remains in advancing the neurobiological framework in ADHD. Consequently, neuroimaging studies have investigated structural, functional, and more recently connectome features in ADHD which may inform the underlying pathophysiology of ADHD presentations. Moreover, growing evidence from these studies may help better reconceptualize ADHD by linking brain-based features with improved clinical care models and treatment outcomes (4).

Research from task and resting-state functional connectivity studies reveal neural differences between the ADHD-C and ADHD-I types with atypical patterns of increased and reduced activation in the cingulo frontal parietal attentional (CFP) and default mode (DMN) networks (5). The regions involved in these networks are of significance to the underlying pathophysiology in ADHD as they are concordant with the clinical deficits associated with the ADHD-I and ADHD-C types, respectively. However, evidence of neuroanatomical differences between the ADHD presentation types remains conflicted, with multiple studies reporting no grey matter volumetric differences (6-10), yet others reporting decreased volumes of the caudate and anterior cingulate cortex (ACC) in ADHD-C relative to ADHD-I type (11), regions associated with nodes of the

DMN in ADHD-I than ADHD-C (12) and also in the hippocampus in ADHD-C relative to ADHD-I type (13). A small number of studies investigating microstructural white matter (WM) properties between the ADHD-C and ADHD-I subtypes, using diffusion tensor imaging (DTI), also yield equivocal findings. DTI is an MRI technique which can provide information on WM microstructure by observing the directionality and coherence of water diffusion (14). This is typically quantified using scalar metrics such as fractional anisotropy (FA), a marker of microstructural architecture; radial diffusivity (RD) to assess axonal myelination; axial diffusivity (AD) as a variable of axonal maturation; and mean diffusivity (MD) as an average of molecular diffusion, independent of directionality (15). Using these *scalar* measures to evaluate the ADHD subtypes, Svatkova, Nestrasil (16) reported increased RD in the forceps minor in ADHD-I relative to ADHD-C (16). Other studies have also reported increased RD bilaterally and AD in mostly left-lateralized fronto-striato-cerebellar regions(17), increased FA and RD in the right thalamus, increased AD in the left postcentral gyrus and right caudate, and increased RD in the left postcentral gyrus and supplementary motor area (18) for ADHD-C relative to ADHD-I. Additionally, studies comparing the ADHD-C type only to controls have observed lower FA values in ADHD-C in regions surrounding the basal ganglia (19, 20). Considering the dearth of studies available in this field, the goal of this study, using both voxelwise analyses of scalar measures and connectomic analyses of tractography measures, was to investigate whether differences in white matter microstructural properties and connectivity, distinguish the two most common ADHD types, ADHD-C and ADHD-I. In this same cohort, we have previously demonstrated differences in graph properties of structural volume covariance networks between the two ADHD types (6). Based on this our goal was to test whether these ADHD types also differed in organization of their brain white matter.

The evolving framework of ADHD pathophysiology is shaped by the application of neuroimaging measures which include the more recent connectomic approach in assessing possible disruptions in brain network connectivity. An example of this paradigm shift can be seen in the proposal of a neurocircuitry based model in ADHD, which incorporates knowledge on the role of inter-regional network organization involving frontal, temporal and parietal regions, from the historical view of dopaminergic regulation and frontal-striatal circuitry deficit (5). Network-based Statistics (NBS) and graph theoretical analysis can be applied to assess structural connectivity, which is represented by anatomical connections formed by WM axonal fiber tracts to understand whether these connections underpin functional network connections. Network topology can be characterized by the application of graph modeling of the structural or functional links characterizing the interregional neuronal connections (21). Only one study to date has compared DTI-based structural connectomic measures between the two ADHD subtypes and found decreased white matter connectivity in ADHD-C compared to ADHD-I, involving mostly right-hemispheric frontal, cingulate and supplementary motor regions (22).

This study utilized DTI data to assess whether structural WM microstructural properties and connectivity may distinguish the ADHD-C and ADHD-I types, and comparatively to neurotypical controls. Firstly, we applied tract-based spatial statistics (TBSS), using DTI derived scalar measures (23) of fractional anisotropy (FA), mean (MD), radial (RD), and axial (AD) diffusivity as indices of water diffusion properties in WM tracts. Secondly, we applied tractography analyses linking 84 brain regions distributed throughout the brain and used network-based Statistics (NBS) to explore connectomic differences and brain network organization, and graph theoretical analysis to capture topographic properties, which may underpin these subtypes.

## Methods

### Participant Characteristics and Study Procedure

Participants were recruited as part of the International Study to Predict Optimized Treatment in ADHD (iSPOT-A) study. A detailed account of the inclusion/exclusion criteria protocols for participant recruitment, diagnostic measures, and procedures for the iSPOT-A study has been previously published (24) and is described briefly below. DTI data collected at Westmead Hospital, Sydney, as part of the baseline MRI data collection for the iSPOT-A study, were available for 37 participants with ADHD (mean = 13.30 ± 2.56; range 8-17 years) and 28 age and gender-matched typically developing controls (mean = 13.09 ± 2.63; range 8-17 years). Data from 2 participants were discarded due to excessive artifacts in the DTI dataset.

Confirmation of ADHD diagnosis (DSM-IV criteria) and subtype (i.e. presentation in DSM-V) was measured by the Mini International Neuropsychiatric Interview (MINI Kid) (25), and the Attention Deficit/Hyperactivity Disorder Rating Scale (ADHD-RS-IV) (26) with symptom severity assessed using the ADHD-RS-IV scores (requires a score of >1 on 6 or more subscale items on the Inattentive and/or Hyperactive/Impulsive subscales) and the Conners’ Parent Rating Scale-Revised: Long Version (CPRS-LV) (24). Of the 37 ADHD participants, 19 met diagnostic criteria for ADHD-C type (mean =13.25 ± 2.53; 4 females), while 18 met diagnostic criteria for the ADHD-I type (mean =13.35 ± 2.65; 4 females). Seven ADHD-C participants and three ADHD-I participants were diagnosed with comorbid oppositional defiant disorder. All ADHD participants were medication-free at the time of testing; 20 were medication naïve; 17 were treatment-experienced with methylphenidate (*n* = 11 ADHD-C; *n* = 5 ADHD-I) and were withdrawn from methylphenidate for at least 5 half-lives. Participants were all fluent in English and had no history of brain injury, any significant medical condition affecting brain function (e.g., epilepsy), or any contraindications for MRI. All participants and/or their guardians provided written informed consent to participate in the research, in accordance with the National Health and Medical Research Council guidelines.

### DTI Image Acquisition and Preprocessing

Magnetic resonance images were acquired using a 3.0 Tesla GE Signa HDx scanner (GE Healthcare, Milwaukee, WI) using an 8-channel head coil. Diffusion tensor images were acquired using a spin-echo DTI-echo planar imaging sequence. Seventy contiguous 2.5mm slices were acquired in an axial orientation with an in-plane resolution of 1.72mm x 1.72mm and a 128 x 128 matrix (repetition time (TR) = 17000 ms; echo time (TE) = 95 ms; fat saturation: on; number of excitations (NEX) = 1; frequency direction: right/left). A baseline image (b = 0) and 42 different diffusion orientations were acquired with a b-value of 1250. Total acquisition time for the DTI protocol was 13 min 36s. Three-dimensional (3-D) T1-weighted magnetic resonance images were also acquired in the sagittal plane using a 3D SPGR sequence (TR = 8.3 ms; TE = 3.2 ms; flip Angle = 11°; TI = 500 ms; NEX = 1; ASSET = 1.5; Frequency direction: S/I). A total of 180 contiguous 1 mm slices were acquired with a 256 × 256 matrix, with an in-plane resolution of 1 mm × 1 mm resulting in isotropic voxels.

Diffusion tensor imaging data pre-processing and analytic methods utilized the Oxford Centre for Functional MRI of the Brain (FMRIB) diffusion toolbox and TBSS software tools as part of the FMRIB Software Library release 4.1.3 (http://www.fmrib.ox.ac.uk/fsl) (23, 27). Raw DTI data were first corrected for head movement and eddy current distortions. Diffusion tensor models were then fitted for each voxel within the brain mask and skeletonized FA, MD, AD, and RD images were generated for each participant. A detailed account of the steps applied prior to running voxel-wise statistical analysis which involves aligning the FA, MD, AD and RD images from each individual using non-linear registration and projecting this onto the mean FA skeleton mask for the cohort, has been published in our previous study (28).

### Tract-Based Spatial Statistical (TBSS) Analysis of DTI Data

Group comparisons were performed for each WM microstructural metric using the Randomise (v2.1) permutation testing software in FSL. We performed three sets of comparisons: (1) ADHD-C versus ADHD-I group, (2) ADHD-C versus controls and (3) ADHD-I versus controls. Permutation testing was performed using 5,000 permutations with the threshold-free cluster enhancement option. We also extracted average DTI values for 46 white matter tract regions defined using the white matter JHU atlas and performed a 3 (group) x 1 (DTI derived FA values) one-way between-groups analysis of variance (ANOVA) (29). We corrected for the number of tracts tested by applying a Bonferroni correction for multiple comparisons, i.e., p < 0.001 (0.05/46).

### Network Analysis of DTI Data

DTI based structural connectome matrices for each individual participant were generated using probabilistic tractography (30). Tractography analysis was performed using each brain region as seed and the remaining region labels as targets. One thousand sample tracts were generated from each voxel within the seed region, and only tracts that reached the target region were retained. The tracts were terminated once they reached a particular target region. The Desikan-Killiany atlas (31) was utilized to define the 84 brain regions using freesurfer (v4.3) (http://surfer.nmr.mgh.harvard.edu/) analysis of the T1 MRI scan. This resulted in an 84 x 84 interregional connectivity matrix of number of probabilistic tracts for each participant.

### Network-Based Statistical (NBS) Analyses

Network-based statistical Analyses (NBS) was then applied to examine regional connectivity differences in the interregional 84 x 84 connectivity matrix between the ADHD-C (n = 19) and ADHD-I (n = 18) types and controls (n = 26). This method has been previously described in detail in Korgaonkar, Fornito (32).

### Graph Theory Analyses

Graph theoretical analyses were performed on the inter-regional 84 x 84 connectivity matrices using the Brain Connectivity Toolbox (http://www.brain-connectivity-toolbox.net/) (33). Global topological properties of the brain were estimated for each individual using: 1) characteristic path length (mean number of connections on the shortest path between any two regions in the network), 2) the clustering coefficient (quantification of the probability that two nodes connected to an index node are also connected to each other). Regional nodal characteristics were measured using the nodal degree (number of connections that a node has with the rest of the network). The matrices were thresholded at a range of network densities in 0.01 steps (0.05 - 0.30) to allow a comparison of network properties between the groups and avoid biases associated with using a single threshold as is typically done. Area under the curve was calculated to examine group differences across the full range of sparsity thresholds, with assessments for both global regional measures performed using a *p* value corrected for a number of global measures, i.e., *p* < 0.025 (.05/2), and for the number of nodes in regional measures, i.e., *p* < 0.0006 (.05/84). This method applied has been previously described in detail (32).

### Correlations between FA values, Network and Clinical Measures

Correlation analyses were performed to measure any associations between: FA values (for the JHU WM atlas tracts) and the global and regional graph network measures, and the dimensional measures of the ADHD-RS scores; total inattention items, total hyperactive/impulsive items and total item scores, while controlling for age and gender.

## Results

Demographic and clinical characteristics for the ADHD-C type, ADHD-I type, and control participants are summarized in Table 1. No significant differences were present between the three groups in terms of age and gender. Medication treatment history, comorbid disorders or the ADHD-RS-IV (inattentive symptom items) sum of items 1-9 did not significantly differ between subtypes. Characteristically, ADHD-C type significantly differed from ADHD-I on the sum of items 10-18 (hyperactive/impulsive symptom items) and total item scores on the ADHD-RS-IV, in concordance with combined type criteria and severity (*p* < .05). Of the ADHD-I type participants, one participant qualified for seven items out of 9 hyperactive/impulsive subscale items, less than four 4 items (*n* = 4), 3 items (*n* = 2), 2 items (*n* = 1) and ≤1 item (*n* = 11).

**Table 1.**
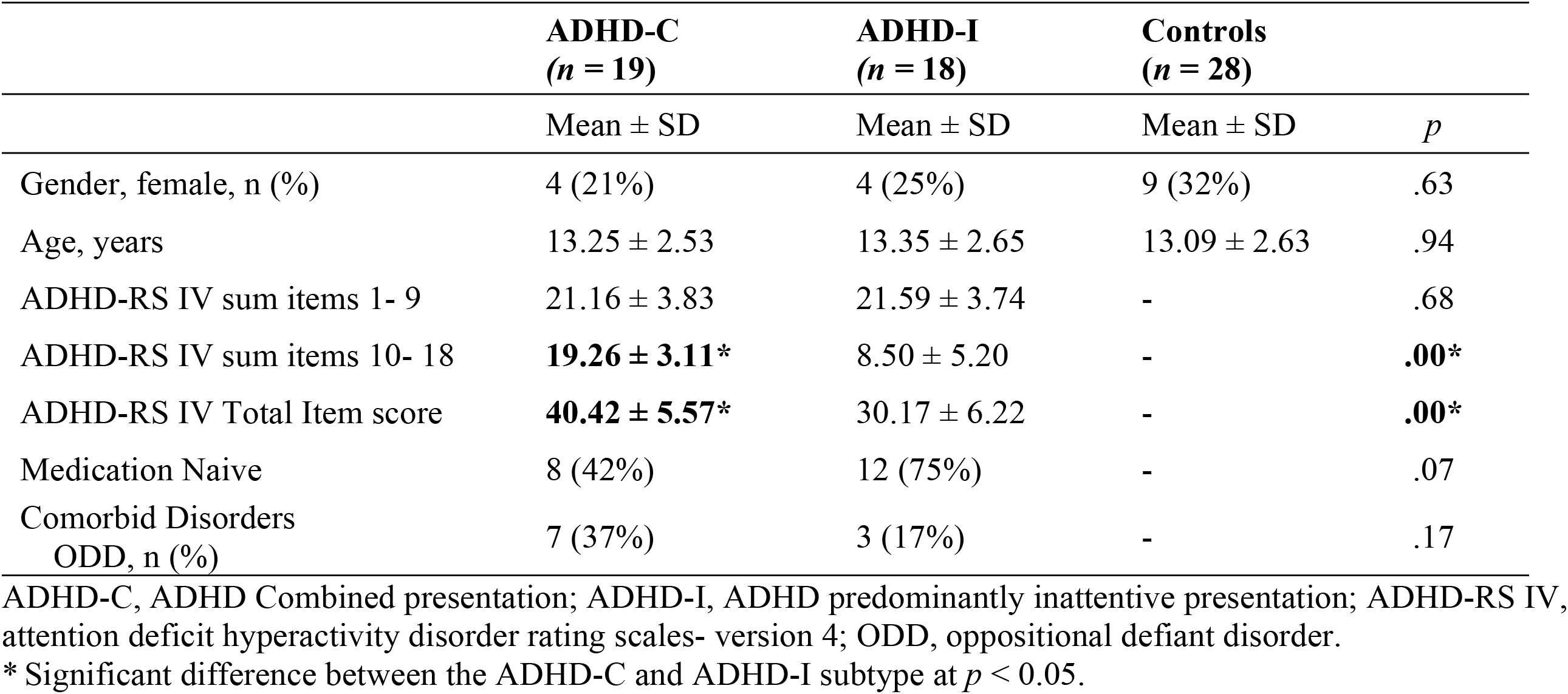
Participant Demographic and Clinical Characteristics.

### Tract-Based Statistical Analyses and Atlas based Region of Interest Analysis

No significant differences were found between the subtypes or relative to controls for any of the DTI measures using both voxel-wise and average values for the 46 defined WM regions. However, findings from the ROI analyses which were significant at the uncorrected level is shown in Fig 1.

**Fig 1.**
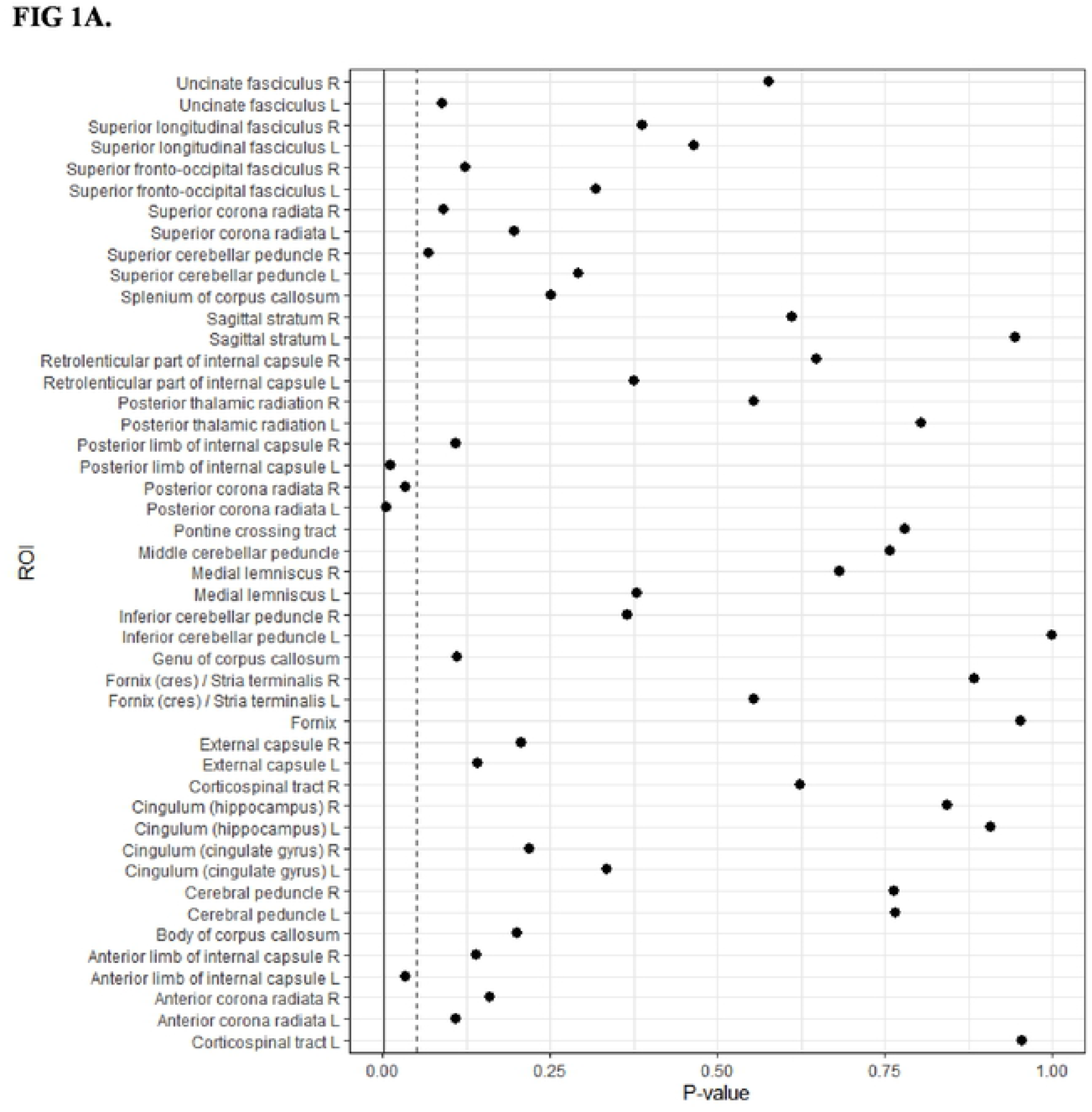
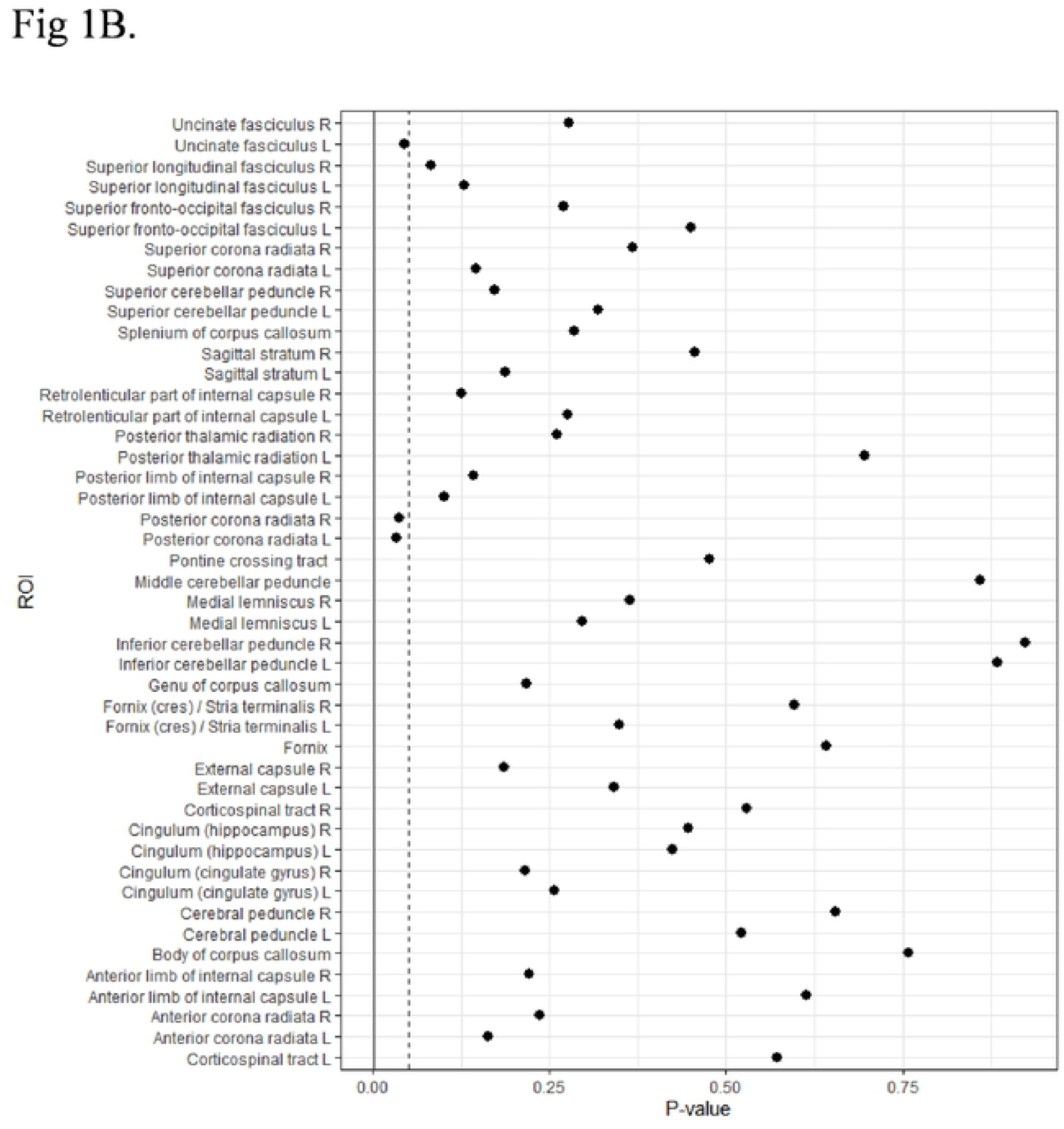
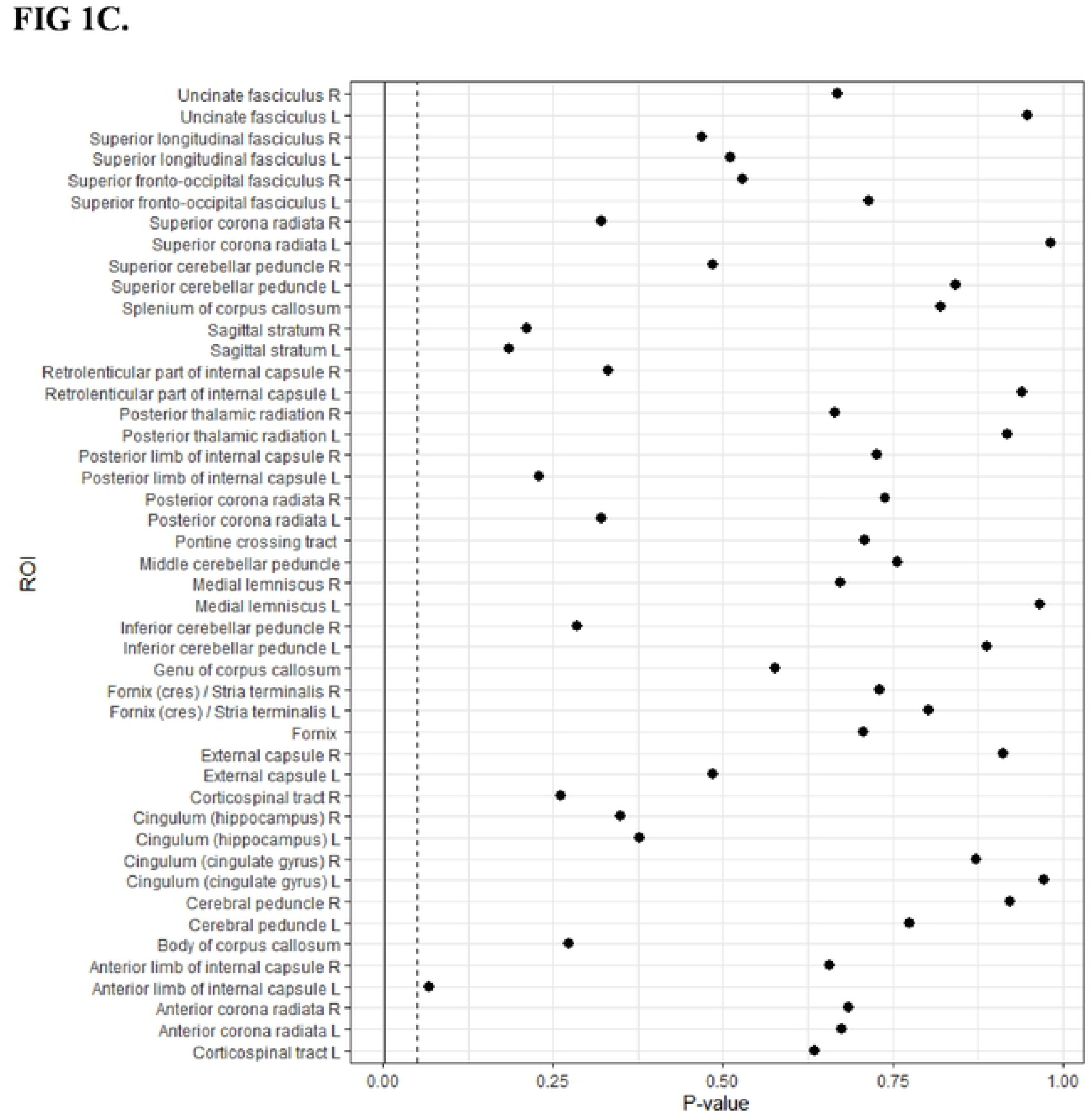
Group comparison of FA values using atlas-based region of interest analysis between the combined (ADHD-C) and inattentive (ADHD-I) subtypes, and controls. Forrest plots showing FA values for 46 defined White Matter Regions using region of interest analysis comparing the combined and inattentive ADHD types (Fig 1A), Combined and controls (Fig 1B), and inattentive relative to controls (Fig 1C). The dotted line represents the .05 p-value threshold, and the solid line represents the Bonferroni corrected p-value threshold.

### Network-Based Statistical Analysis

NBS analysis also did not identify any inter-regional connectivity differences between the ADHD-C and ADHD-I types or when compared to controls after correcting for multiple comparisons.

### Graph Theoretical Analysis

Neither of the global topological measures: clustering coefficient or global characteristic path length, showed significant differences between the ADHD-C and ADHD-I groups or compared to controls. Group comparison of the regional nodal degree revealed a single significant finding between the ADHD-C and control group, (*p* < .05 corrected for multiple comparisons) in the right insula (corrected *p* = .029; ADHD-C < Controls). There were no significant differences found between the two ADHD subtypes, or the ADHD-I type compared to controls. Regional nodal degree measures which were significant but did not survive after correcting for multiple comparisons are illustrated in Fig 2A, 2B and 2C.

**Fig 2.**
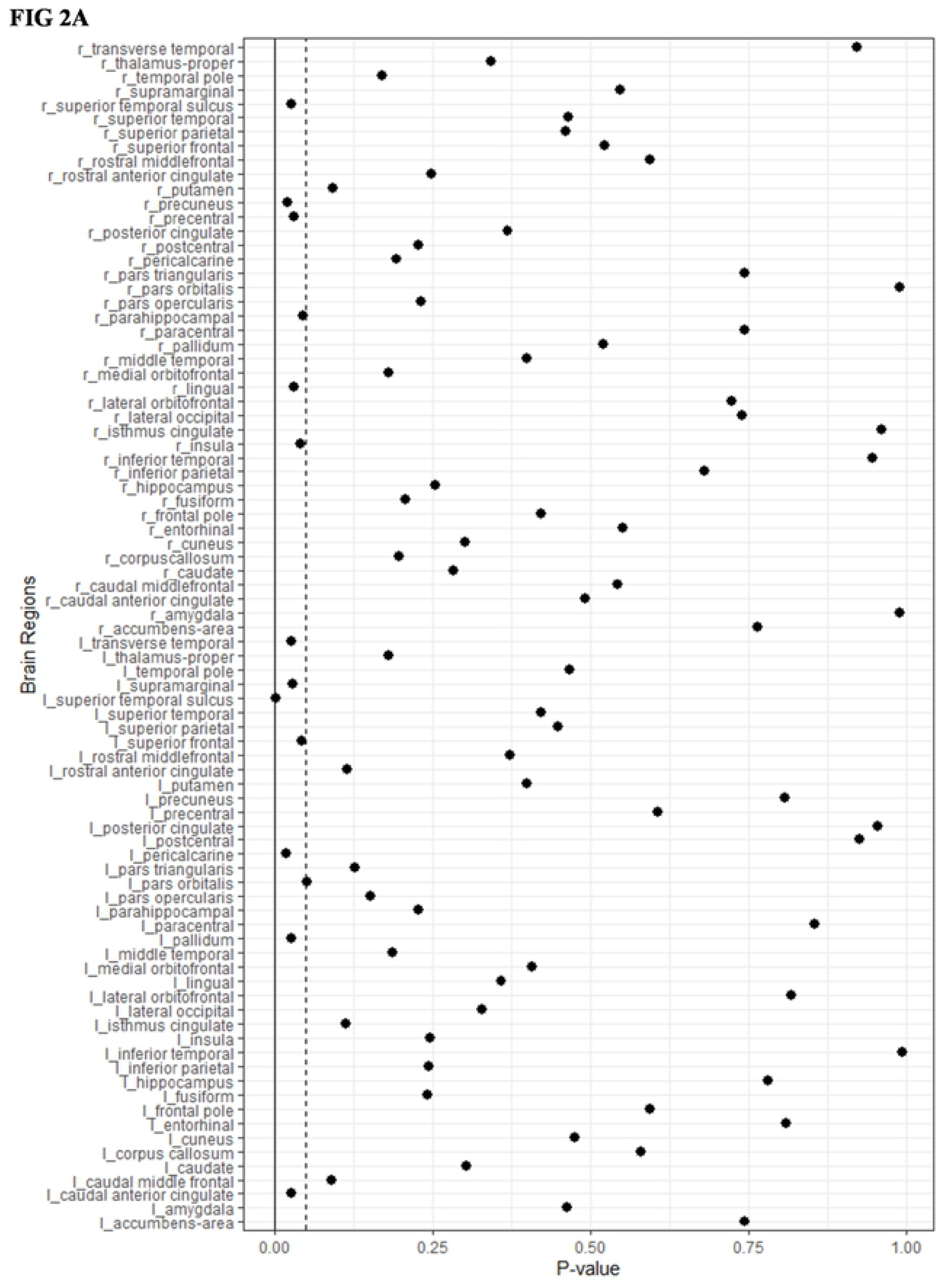
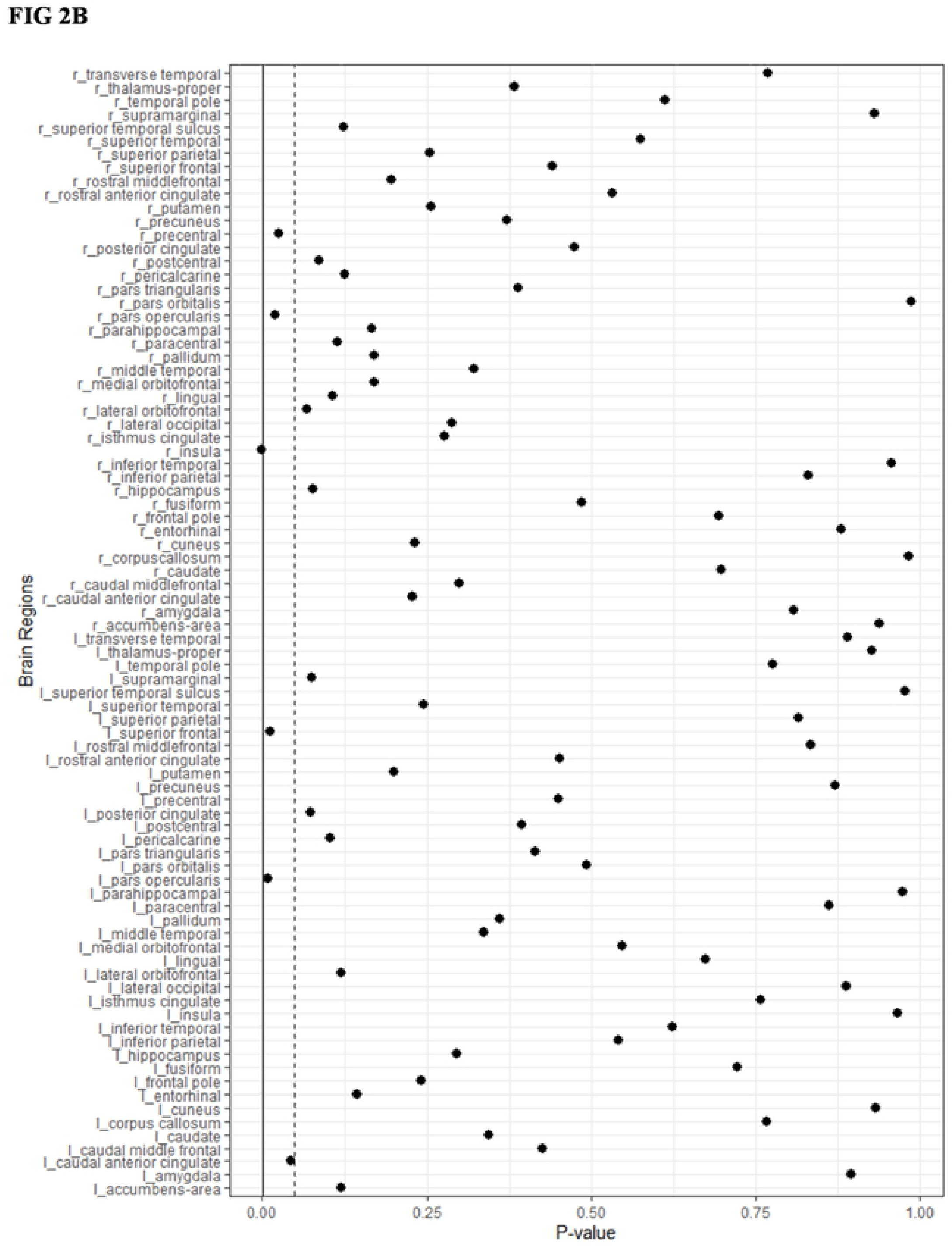
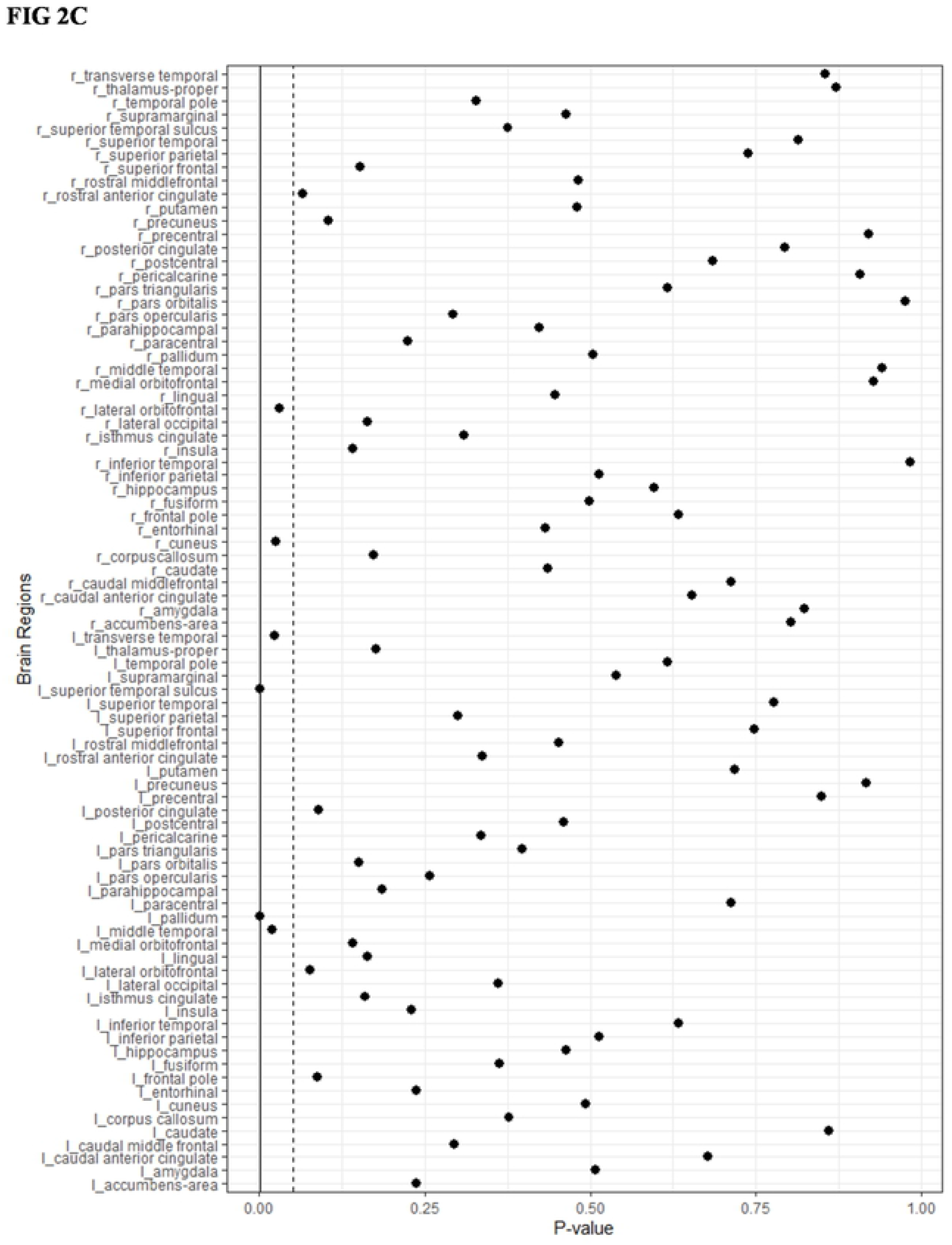
Group comparison of regional nodal degree measures using graph theoretical analysis between the combined (ADHD-C) and inattentive (ADHD-I) subtypes, and controls. Forrest plots showing p-values for 84 brain regions comparing the combined and inattentive ADHD types (Fig 2A), Combined and controls (Fig 2B), and inattentive relative to controls (Fig 2C). The dotted line represents the .05 p-value threshold, and the solid line represents the Bonferroni corrected p-value threshold.

### Correlations between the ROI, Network and Clinical Measures

While no significant associations were found following Bonferroni correction, associations were observed at the uncorrected level between the sum of items 1-9 and FA in the left anterior limb of the internal capsule; *p* = 0.039, both the sum of items 10-18 and total item score with FA in the right superior cerebellar peduncle at *p* = 0.032 and *p* = 0.049, respectively. None of the global or regional network measures were correlated with the clinical rating measures of the ADHD-RS-IV. Associations observed between regional nodal degree measures and clinical measures at an uncorrected threshold have been summarized in Table S1, S2 and S3 in the supplementary material.

## Discussion

This study investigated whether structural WM microstructural properties and inter-regional white matter connectivity distinguished the combined (ADHD-C) and inattentive (ADHD-I) DSM-V presentations of ADHD. TBSS analysis of FA, AD, MD, and RD values, ROI analysis of FA values for 46 white matter tracts defined by the JHU WM atlas and NBS analyses of inter-regional connectivity of 84 cortical-subcortical brain regions revealed no significant differences between the subtypes or compared to controls. Of all the white matter measures, only graph properties of nodal degree for the insula were found to significantly differ in the ADHD-C type relative to controls after controlling for multiple comparisons. Studies of WM microstructural properties between the subtypes of ADHD are limited. In line with the only two previous TBSS studies (16, 17) available in the literature, our findings did not find differences in FA, between the two ADHD subtypes. However, both the Svatkova (16) and Ercan (17) studies did observe differences in RD, MD, and AD between the two ADHD types. An earlier study (18) demonstrated FA, RD, and AD differences between the two subtypes, although the analytical approach employed in that study is no longer accepted in current DTI studies which favor TBSS, as a sophisticated model for measures of DTI scalar values (34). While we did not find any significant white matter tractography differences between the combined and inattentive subtypes of ADHD, only one other study exists that has reported on white matter connectivity differences in the frontal, cingulate and supplementary motor regions between patients with ADHD-C and ADHD-I type using NBS (22). As such, further research is warranted to determine differences in WM structural integrity between the hyperactive/impulsive subtype relative to the other subtypes.

Another possible reason for null findings in our study may be due to a number of treatment-experienced participants in our study compared to the previous subtype-based studies, which recruited participants who were treatment-naive or with treatment history < 6 months (17, 18, 22). Svatkova, Nestrasil (16) had mostly treatment-experienced participants (n = 21, 63%) without washout prior to scanning. Participants in our study who were treatment-experienced (n = 17, 46%) commenced scanning procedures after entering a washout period of 7 days. Interestingly, previous subtype inclusive studies which compared ADHD participants to controls and found significant differences also had a majority of treatment naïve participants (35-38). However, it is important to note that these previous studies are not comparable with our study in terms of methodological approaches. Furthermore, a recent DTI study investigated a medication naïve ADHD group of children and adults and found no differences to controls in the child group but reported significant findings in the treatment naïve adult group (39). Nevertheless, it may be likely that white matter differences in ADHD may be influenced by stimulant treatment effects but may also be due to neurodevelopmental periods in line with the proposed maturational lag theory in ADHD (40).

Consistent with the two previous subtype comparison studies utilizing TBSS, we found no differences in FA values (16, 17) or AD and MD values, distinguishing the two types (16). Notably, while both studies employed cohorts that were similar to our study in terms of clinical characteristics and sample size, the Ercan, Suren (17) study included a treatment naïve sample. Interestingly, increased FA values were found relative to controls, in the bilateral cingulum bundle in ADHD-C, and the anterior thalamic radiation, inferior longitudinal fasciculus and left corticospinal tract involving regions linked to fronto-striatal-thalamic circuits in ADHD-I (16). Increased RD in the forceps minor in ADHD-I relative to ADHD-C (16) and increased RD bilaterally and increased AD in brain regions mostly on the left side linked to fronto-striato-cerebellar regions in ADHD-C relative to ADHD-I (17) have also been reported. In contrast, also using a TBSS approach, Rossi et al. (41) measured FA values for 20 fiber tracts in ADHD-I children compared to controls and found no significant difference between the groups.

Additionally, we performed a region of interest (ROI) analysis and did not find significant differences in FA values between the two types. Despite the absence of significant differences between groups at the corrected thresholds, increased FA values in ADHD-C compared to ADHD-I and controls were observed at uncorrected thresholds. That is, the bilateral posterior corona radiata characterized the ADHD-C type from ADHD-I and controls, the left posterior limb of internal capsule and left anterior limb of internal capsule in ADHD-C relative to ADHD-I, and the left uncinate fasciculus in ADHD-C compared to controls. Interestingly, similar findings from a previous study comparing an ADHD group reported reduced FA values in the ADHD groups in the posterior corona radiata, the uncinate fasciculus with reduced MD in the posterior limb of internal capsule, relative to controls (42).

In our study, we also used tractography to evaluate white matter connectomic differences. Our study did not find WM connectivity differences between the ADHD-C and ADHD-I subtypes or compared to controls. However, previous findings from Hong, Zalesky (22), the only other study evaluating connectome differences between the subtypes, observed decreased WM connectivity differences in ADHD-C relative to ADHD-I connecting 17 right hemispheric brain regions including the superior frontal gyrus, anterior cingulate gyrus, and supplementary motor areas. Additionally, they found significant negative correlations between FA values and continuous performance task scores to differentiate ADHD-C and ADHD-I. Also, we did not find any significant differences between the two ADHD subtypes for global topological characteristics or regional nodal graph measures. However, a single significant finding was observed which showed reduced nodal degree involving the right insula in the ADHD-C group when compared to controls. Whilst speculative, similar findings involving the insula have been noted in previous structural (43-45) and functional connectivity (46, 47) studies in ADHD and warrant further research. The insula is considered part of the salience network which interacts with the default mode network. Reduced degree is this region may account for attentional processing and goal-directed action deficits and tied to mechanisms of executive functioning and sensorimotor dysregulation associated with ADHD-C type (48). Furthermore, this finding may have implications for attentional and inhibitory emotional processing in ADHD (45). At the uncorrected thresholds, regional network measures revealed some interesting results; with increased nodal degree in regions involving the occipital temporal, and parietal lobes in ADHD-I relative to ADHD-C and controls and increased nodal degree in regions of the frontal lobe and the basal ganglia in ADHD-C compared to ADHD-I and controls. These findings are consistent with the Hong, Zalesky (22) study that also revealed decreased connectivity in ADHD-C, relative to ADHD-I, in frontal regions, which are involved in regulating inhibitory response and motor activity, both of which are linked to the ADHD-C type.

Although the availability of DTI studies which specifically examine subtype differences are sparse, significant findings have been reported, albeit equivocal. Undoubtedly, further studies are warranted to establish structural connectivity alterations in ADHD types, however, emerging evidence from DTI studies examining an all-inclusive ADHD group with controls have reported significant findings. For example, studies investigating basal ganglia and thalamic connectivity have found lower FA values in ADHD-C relative to controls (19, 20). Fall and colleagues (49) indirectly support these findings reporting mean reaction times correlated with MD values in the striatum and thalamus in ADHD-C compared to controls. Interestingly, these results are consistent with studies using structural gray matter and resting state functional connectivity, reported in a recent review (50) which have shown disruptions in the motor network in ADHD-C. Whether these structural connectivity disruptions or the absence of WM abnormalities attribute to functional deficits associated with the subtypes of ADHD, remain unconfirmed. And so, while the present study focused on whether hard-wired networks may shed insight that distinguishes the subtypes, further exploration of the functional dynamics of brain network organization may potentially provide greater insight toward the underlying neurobiology of the ADHD subtypes. Importantly, the rationale for the DSM-V based diagnostic criteria adopting ‘presentation type’ in place of ‘subtypes’ was to try to convey the somewhat transient nature to these classifications and how these clinical symptoms evolve throughout development (50, 51). Perhaps this is another potential reason for the null findings as ADHD presentation types are not stable across time, and thus they may be better linked with functional measures.

### Limitations

There were several limitations to the study. It is quite likely that the lack of significant findings in our study is due to the small available sample size. This limits the statistical power of our results and thus may explain why we were not able to detect differences between groups after appropriate correcting for multiple comparisons. We have reported uncorrected findings in the supplementary section and replication studies with larger sample sizes utilizing these measures are warranted to further explore possible differences in microstructural integrity between the ADHD types. In a previous study utilizing the same cohort, we did observe differences in structural covariance between the two ADHD subtypes. It may be that interregional structural covariance relates better to functional networks and it is likely that functional differences are more pronounced between the ADHD types (6, 52). It is, however, important to note that some of the previous studies have reported significant white matter differences with similar sample sizes. It is likely these results could be cohort-specific, stressing the need for replication in larger cohorts. We applied standard established analytical methods and hence our null findings are important to consider in the context of the paucity of DTI studies comparing ADHD presentations.

Our study examined the combined and inattentive type only, as data for the predominantly hyperactive-impulsive type was not available, thereby analysis between all three types was not possible. Medication effects and stimulant treatment history may bias the findings from this study; however, this confounding effect is difficult to eliminate. Although, recent ADHD DTI research studies which investigated possible medication effects on WM structure and connectivity produced inconclusive findings (39). Changes in FA may be influenced by variable factors such as familial vulnerability, ADHD symptom count, and neurodevelopmental periods as it is well established that FA increases from 10–12 years up to 40 years of age in all WM tracts (53, 54). Consequently, these factors are not uniformly accounted for in ADHD research and may limit the generalizability of the results of our study. While this study did match participants for age and gender, analyses could not be explored by developmental periods to explore possible FA differences due to the small sample size. Finally, DTI lacks any information on functional connectivity of the brain and as such replication of these findings will follow in another study which will involve resting-state fMRI to determine whether similarities are found in functional interregional connections between the two subtypes.

## Conclusion

In summation, we did not find white matter microstructural properties or network connectivity to differentiate the two subtypes from each other or relative to controls. However, we did find the ADHD-C group to differ in regional nodal degree in the insula relative to controls. While the overall results of this study may be inconsistent with the findings of previous ADHD subtype DTI studies, the absence of evidence for differences in WM microstructural properties relative to FA values between the two subtypes is in line with previous research. To the best of our knowledge, this is the first study to utilize both voxel-wise analysis of scalar white matter measures in addition to tractography incorporating network and graph theoretical measures to examine differences between these two ADHD subtypes.

## Acknowledgments

The iSPOT-A trial was sponsored by Brain Resource Company Operations Pty Ltd. We acknowledge the NHMRC funded Project Grant (APP1008080) awarded to Korgaonkar, Grieve & Williams for the provision of some of the control data. Prof Leanne Williams was the academic Principal Investigator for iSPOT-A (2009-2013), and Prof Simon Clarke was the Principal Investigator for the iSPOT-A Sydney site. Dr. Kristi Griffiths is supported by an NHMRC Early Career Researcher Fellowship (GNT1122842). Dr. Korgaonkar is supported by an NHMRC Career Development Fellowship (APP1090148). We thank Dr. Lavier Gomes and Ms. Sheryl Foster and the Department of Radiology at Westmead Hospital for their substantial contributions to magnetic resonance imaging data acquisition. We thank Tracey Tsang who served as the iSPOT-A trial coordinator, along with Sariah Hobby, Yennie Hyunh and Jodie Logan who assisted in data acquisition. We also thank the individuals who gave their time to participate in the study.

## Supporting Information

**S1, S2, S3 Tables. Correlations between the ADHD-RS IV scores and diffusion metrics of children and adolescents with adhd combined and adhd inattentive**

**S1 Table. Correlations between the global network measures and the ADHD-RS IV scores**.

**S2 Table. Correlations between averaged FA values of the 46 white matter tracts and the ADHD-RS IV scores**.

**S3 Table. Correlations between regional nodal degree of the 84 white matter regions and the ADHD-RS IV scores**.

